# Breathlessness in a virtual world: An experimental paradigm testing how discrepancy between VR visual gradients and pedal resistance during stationary cycling affects breathlessness perception

**DOI:** 10.1101/2022.06.16.496494

**Authors:** Sarah L. Finnegan, David J. Dearlove, Peter Morris, Daniel Freeman, Martin Sergeant, Stephen Taylor, Kyle T.S. Pattinson

## Abstract

**Introduction:** The sensation of breathlessness is often attributed to perturbations in cardio-pulmonary physiology, leading to changes in afferent signals. New evidence suggests that these signals are interpreted in the light of prior “expectations”. A misalignment between afferent signals and expectations may underly unexplained breathlessness. Using a novel immersive virtual reality (VR) exercise paradigm, we investigated whether manipulating an individual’s expectation of effort (determined by a virtual hill gradient) may alter their perception of breathlessness, independent from actual effort (the physical effort of cycling).

**Methods:** Nineteen healthy volunteers completed a single experimental session where they exercised on a cycle ergometer while wearing a VR headset. We created an immersive virtual cycle ride where participants climbed up 100 m hills with virtual gradients of 4%, 6%, 8%, 10% and 12%. Each virtual hill gradient was completed twice: once with a 4% cycling ergometer resistance and once with a 6% resistance, allowing us to dissociate expected effort (virtual hill gradient) from actual effort (physical effort of pedalling). At the end of each hill, participants reported their perceived breathlessness. Linear mixed effects models were used to examine the independent contribution of actual effort and expected effort to ratings of breathlessness (0-10 scale).

**Results:** Expectation of effort (effect estimate ± std. error, 0.63 ± 0.11, *p*<0.001) and actual effort (0.81 ± 0.21, *p*<0.001) independently explained subjective ratings of breathlessness, with comparable contributions of 19% and 18%, respectively. Additionally, we found that effort expectation accounted for 6% of participants’ physical effort of pedalling and was a significant, independent predictor (0.09 ± 0.03; *p*=0.001).

**Conclusions:** An individuals’ expectation of effort is equally important for forming perceptions of breathlessness as the actual effort required to cycle. A new VR paradigm enables this to be experimentally studied and could be used to re-align breathlessness and enhance training programmes.

## Introduction

Breathlessness is a complex perception, in which the brain is now recognised as an active contributor. Rather than passively transmitting sets of afferent signals, new evidence suggests that the brain interprets incoming signals based on a set of held “expectations” [1–3]. Because perceptions, including those of breathlessness are generated in the brain, a disconnect between the lungs and perception can be explained as a mismatch between the senses [1, 4, 5]. Such a disconnect may explain why breathlessness is often difficult to treat, with many people, such as those living with chronic lung or cardiac disease remaining symptomatic despite maximal medical therapy [6–8]. For these people, the sensation of breathlessness may not match the physical status of the cardiorespiratory system. Indeed, sensory mismatches resulting in breathlessness can be generated experimentally via engendered expectations to placebo and nocebo cues [9]. Conversely, treatments drawing on exposure based cognitive therapies, which focus on changing sensory and emotional expectations, appear to be most successful in providing symptomatic relief from chronic breathlessness [10, 11].

Current evidence highlights that an individual’s expectations about breathlessness and exercise may influence their subjective experiences. For example, an individual who realises they have forgotten their inhaler may suddenly experience feelings of shortness of breath. However, given that breathing contains both conscious and unconscious elements, is emotionally complex and difficult to articulate, quantifying this relationship is challenging.

Recent work has built on well-established embodiment illusions, such as the rubber-hand illusion [12], in which participants “feel” the rubber hand as their own to show how visual and somatosensory cues can be used to “trick” perceptions of bodily status. Using virtual avatars, manipulating the avatar’s respiratory phase directly altered feelings of participant self-location, breathing agency and tidal volume [13]. The affective impact of breathlessness was also attenuated by the presence of a virtual avatars breathing, particularly when asynchronous with a participants own breathing [14]. Manipulating this sense of agency is not restricted to passive situations or to external cues. While exercising under hypnosis, ratings of perceived exertion have been shown to reflect the suggested, rather than actual work effort [15]. Similarly, during imagined hand-grip exercises, individuals with high hypnotizability perceived their rates of exertion as higher than those with low hypnotizability [16].

One key aspect that drives expectation is sensory immersion, which now, rather than relying on hypnosis or carefully contrived illusions, is immediately available under full control via Virtual Reality (VR) technology. As a result, VR technology is emerging as a powerful tool within healthcare and research settings, allowing therapeutically relevant situations to be created, repeated and manipulated with a consistency impossible to create in real life [17, 18]. The situations created by VR are immediately available (and readily ended if participants become uncomfortable), facilitating patient access and engagement in a safe space. Meanwhile, researchers can create “real-world” environments whilst maintaining carefully controlled experimental parameters. Thus, VR has the potential to disentangle how prior expectations contribute to breathlessness.

Virtual reality has already been used therapeutically to reduce complex regional pain [17], improve the acceptance and embodiment of prosthetic limbs [18] and facilitate recovery post-stroke [19]. However, VR interventions for chronic breathlessness are in their infancy. Gamification of physical exercise has been positively received [19–21], with participants reporting higher enjoyment and self-efficacy compared to traditional static cycling. A recent systematic review which found moderate effects of VR-exercise (including full body work out, jogging and balance training) on breathlessness, and weak effects on heart rate and oxygen saturation over traditional exercise, highlighted the limited number of studies and lack of standardization of methodology. Furthermore, none of the virtual gaming interventions were immersive, i.e. using a reality headset. Therefore, it remains unknown whether perceptions of breathlessness can be independently manipulated via a VR exercise paradigm.

Here we used a novel immersive VR exercise paradigm to determine whether breathlessness could be manipulated independently of physical effort by modulating an individual’s expectation of effort.

## Methods and Materials

### Participants

Nineteen healthy adult participants (10 female, median age 20 years; range 19-41 years) were recruited to the study. Written informed consent was obtained from all participants prior to enrolment. The study was granted approval by University of Oxford, Medical Sciences Interdivisional Research Ethics Committee (Ref: R68447/RE001). Study inclusion criteria were: adults aged 18-65, a willingness and ability to provide informed consent, and a willingness and ability to ride a bicycle. Exclusion criteria were: inadequate understanding of verbal and written English; significant cardiac, neurological, psychiatric (including depression under tertiary care) or metabolic disease (including insulin-controlled diabetes); history of cardiac tachyarrhythmia or of prescription/non-prescription drug dependency (including alcoholism).

### Study visit protocol

Following telephone screening, participants were invited to attend a single one-hour session at the Department of Physiology, Anatomy and Genetics, University of Oxford.

#### Self-report questionnaires

To assess whether baseline measures of anxiety influenced either perceived breathlessness or modulated the interaction between VR hill gradient and physical effort, participants completed the state component of the State-Trait Anxiety Inventory (STAI) [20] prior to commencing exercise. To assess levels of immersion within the virtual environment we collected scores on the Presence Questionnaire [21]. Questionnaires were scored according to their respective manuals.

#### Anthropometric measurements

Participant height and weight were recorded upon arrival to the laboratory.

#### Cycling protocol

Exercise was performed on a bicycle attached to a Tacx Flow Smart turbo trainer (Garmin Ltd., US) that recorded power output and communicated directly with the VR software (Figure 1A). The turbo trainer was calibrated prior to each exercise session and tyre pressure was maintained at 90 Psi. Participants adjusted the bicycle saddle height according to personal preference and ambient conditions were controlled.

**Figure 1.**
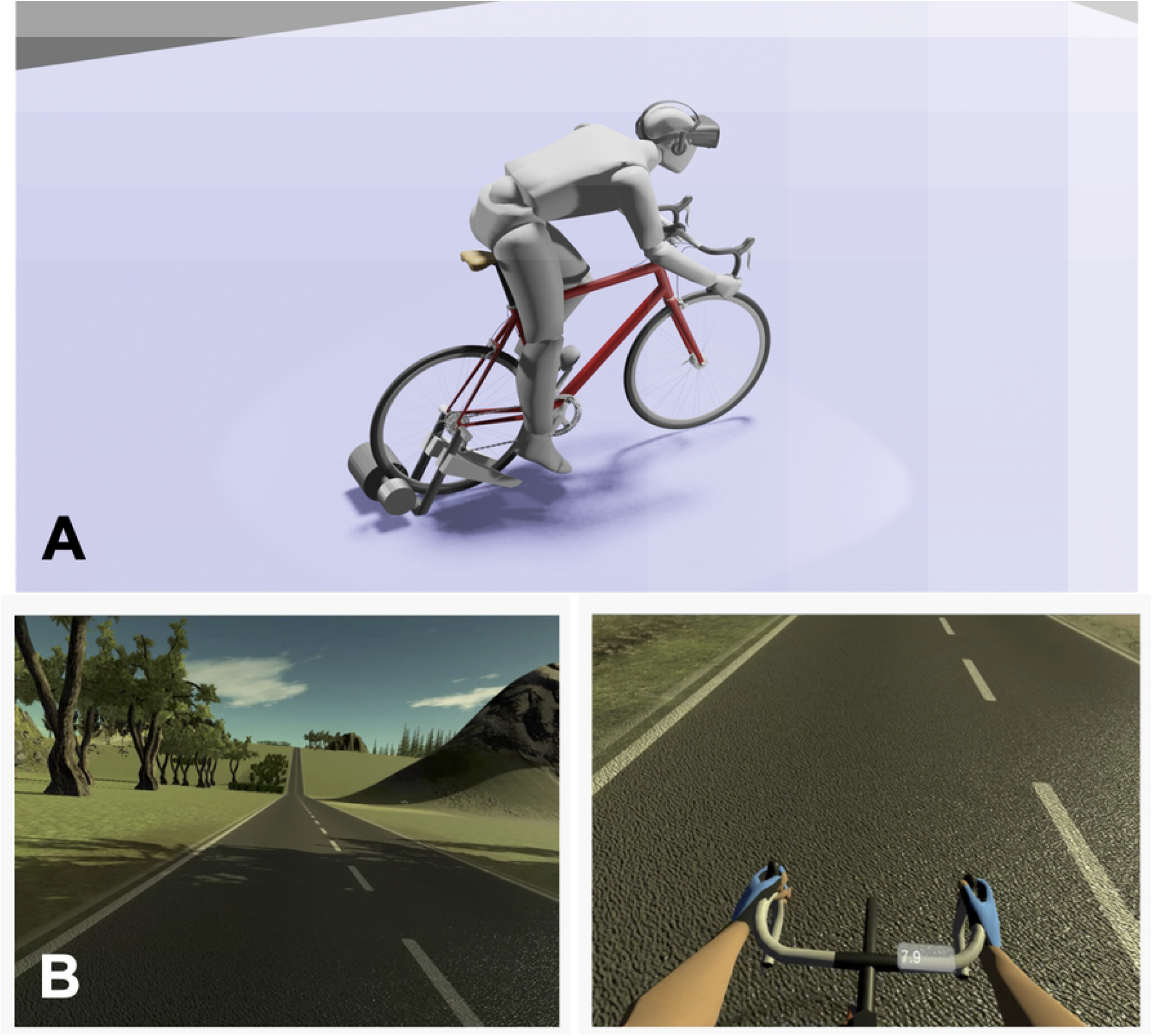
A. Image of the bicycle ergometer set-up and B. (top) the VR environment (bottom) virtual hands and speedometer.

Participants completed two bouts of cycling in a virtual world with a rolling terrain (Figure 1b), where the physical effort of pedalling (slope resistance value between 0 to 100, with 0 representing a flat surface and 100 representing a 6% gradient - the maximum gradient simulation on the Tacx Flow Smart turbo trainer) was independently manipulated from the observed VR hill gradient. The software was written in Unity (version 2019.2.17), using the Unity packages VRTK (version 3.3.0) for integration with VR and Advanced ANT+ (version 1.041) for communication with the turbo trainer (or any fitness equipment using the ANT+ protocol). In addition, an Arduino UNO microcontroller was integrated into the system, to allow analog/digital inputs. The code is available on request at https://github.com/Taylor-CCB-Group/BreathlessVR. The virtual world was viewed on an Oculus Rift headset.

The first exercise bout was a familiarisation session and served to ‘anchor’ participants perceptions of breathlessness. Participants rested for an average of 4 minutes 40 seconds before starting the second exercise bout. Only data from the second exercise bout were included in the final analyses. In each exercise bout, participants warmed-up for 300 m before completing an undulating course containing 12, 100 m long hills with observed gradients of 4%, 6%, 8%, 10% or 12% (Figure 2 and Supplemental Table 1). Participants completed each hill gradient twice, once with a slope resistance of 4% and once with a slope resistance of 6%. In so doing, we sought to dissociate actual effort (i.e., the physical effort of pedalling) from participants’ expectation of effort (i.e., the virtual hill gradient). In between each work block, participants completed a 30 m recovery ride on a flat surface (0% gradient) with a slope resistance value of 1%. After each 100 m hill, a virtual prompt appeared asking participants to report how breathless they felt on a scale of 1-10 (1 being not at all breathless and 10 being maximum breathlessness). Participants’ verbal report was recorded automatically by the computer, and this was verified by a study investigator. The block order for the second exercise bout. Participants were counter-balanced in receiving block order one or two first. Participants cycled at a self-selected pace.

**Figure 2.**
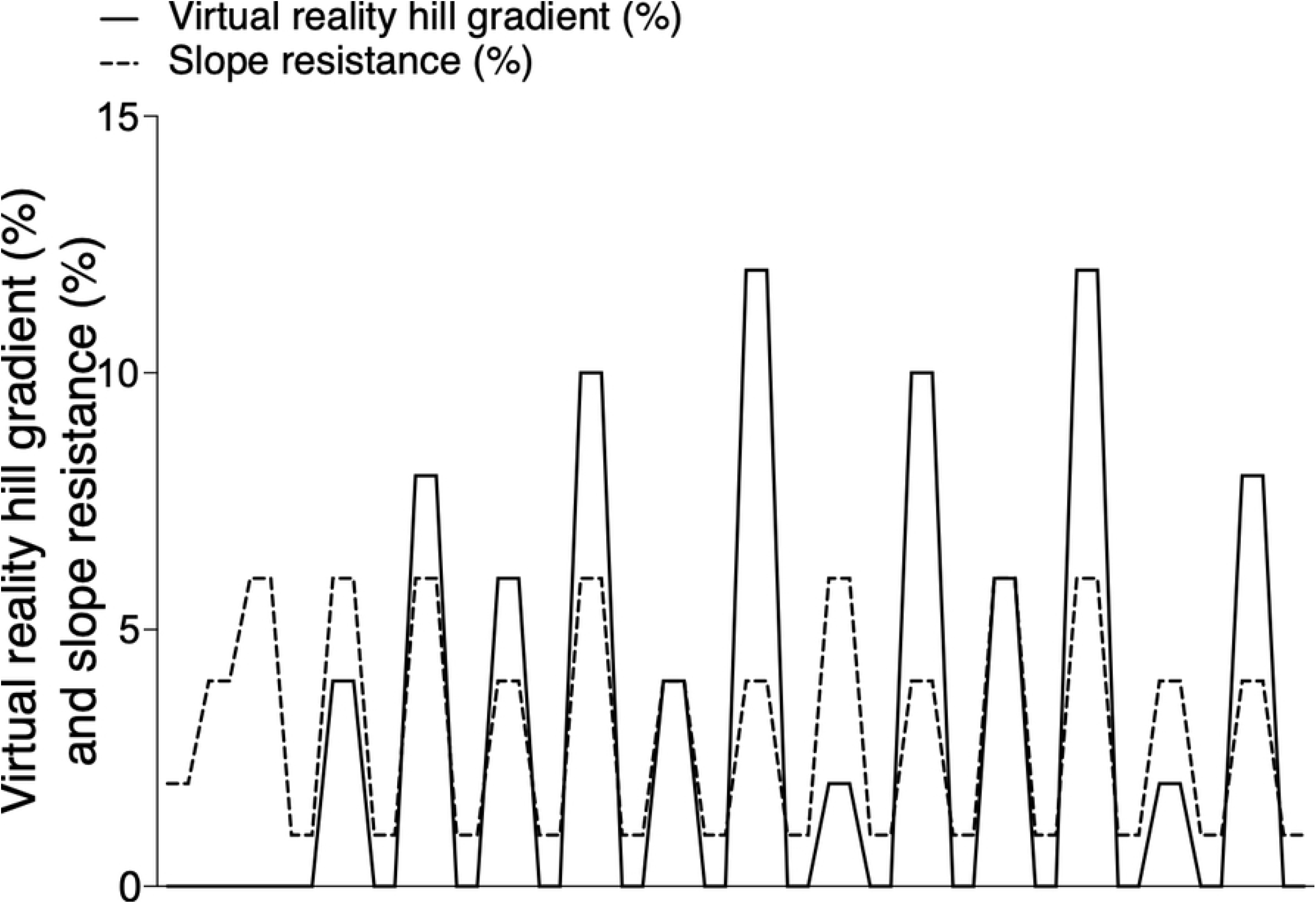
Illustration of exercise protocol where slope resistance (cycling ergometer resistance) was dissociated from the observed VR hill gradient.

#### Analysis

All variables were centred and scaled prior to analysis. To assess whether slope resistance (4% and 6%) affected participants’ physical effort (W), a 2-way repeated-measures ANOVA was performed. To determine the effect of actual effort and effort expectations on breathlessness, and to determine the effect of effort expectation on actual effort, linear mixed-effects models were created in RStudio (Version 1.3.1093) using the lme4 package [22]. For both models, a random intercept was fitted for each participant. Final models (combinations of predictor variables) were selected by Akaike Information Criterion (AIC) by backwards elimination using the stats package [23]. Full model summaries and terms are presented in Supplementary materials.

## Results

### Participant characteristics

Nineteen participants completed the trial and were included in final analyses (see Table 1 for participant characteristics). Participants were on average below clinical thresholds for anxiety, typically defined as a score of 40 or more [24] (Table 1).

**Table 1.** Demographic and questionnaire data collected for 19 participants. BMI – body mass index, STAIT – state anxiety questionnaire.

| N=19 | Score |
| --- | --- |
| Age (median years/range) | 20 (19-41) |
| BMI (kg.m<sup>-2</sup> ± SD) | 22 ± 2 |
| Gender – Female | 11 |
| STAIT (IQR) | 25 (10) |

### Exercise time to completion

The time taken to complete exercise blocks 1 and 2 was comparable (*p*>0.05; Table 2).

**Table 2.** Average cycling and rest durations across 19 participants.

| Block 1 (std) | Rest (std) | Block 2 (std) | Average Exercise (std) |
| --- | --- | --- | --- |
| 7 minutes 12 seconds<br>±<br>1 minute | 4 minutes and 40 seconds<br>±<br>1 minute 48 seconds | 6 minutes 39 seconds<br>±<br>1 minute 10 seconds | 6 minutes 55 seconds<br>±<br>1 minute 7 seconds |

### Association between slope resistance and power

To test the hypothesis that participants worked harder during the 6% vs. 4% slope resistance intervals, a 2-way ANOVA was performed where power (W) was explained as a function of slope resistance, VR hill gradient and their interaction (see Supplementary Table 2 for full ANOVA results). There was a significant main effect of slope resistance (*p*<0.001; Figure 3), demonstrating that participants produced more power in the 6% vs. 4% intervals. This relationship was consistent across all levels of VR hill gradient, as evidenced by a non-significant interaction effect (*p*>0.05).

**Figure 3.**
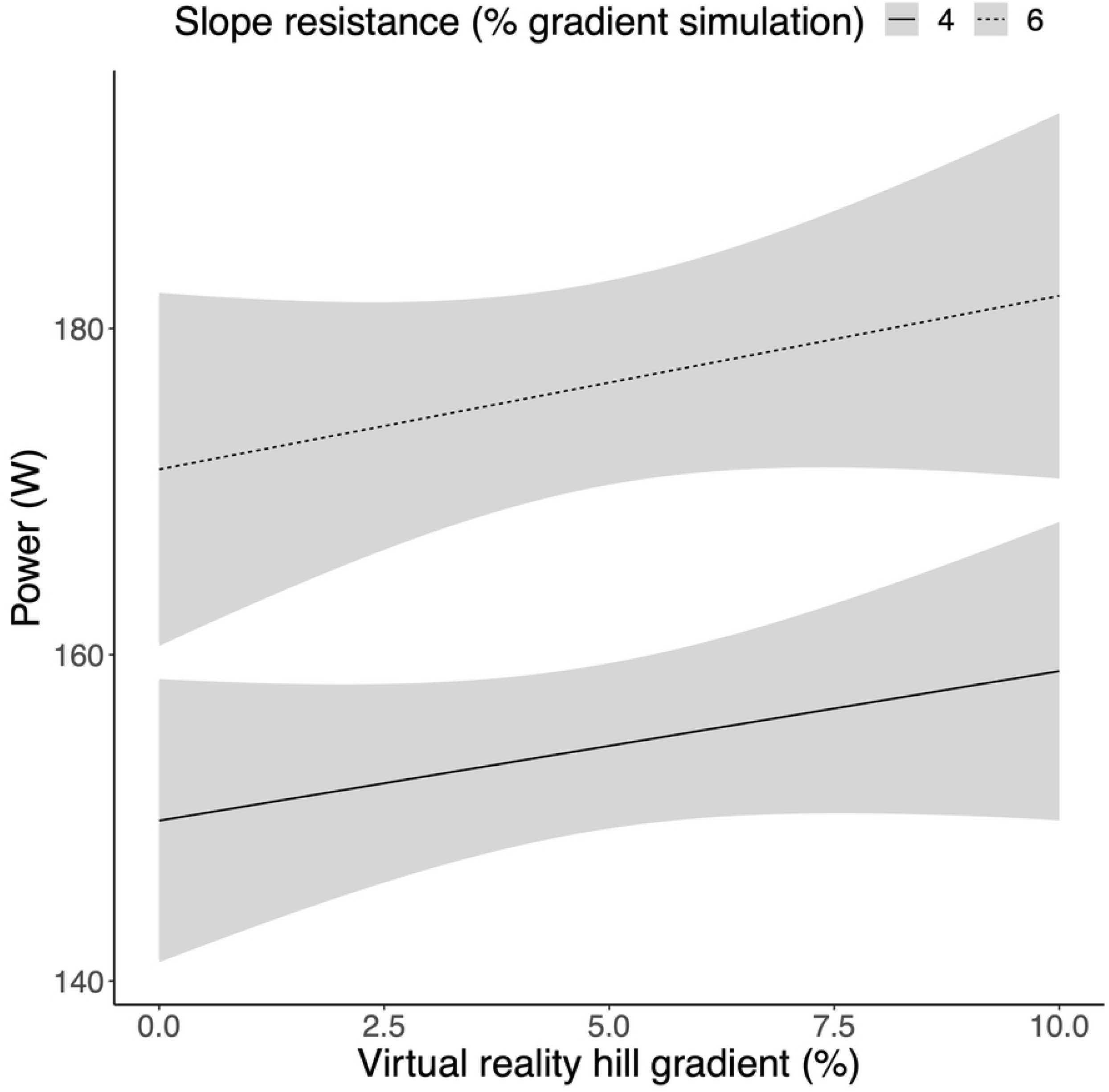
Slope resistance is a significant predictor of power. Mean power (W) was greater across all levels of VR hill gradient in the 6% vs. 4% slope resistance intervals. Shaded areas represent 95% CIs.

### Virtual reality effort is an independent predictor of breathlessness

To determine whether an individual’s expectation of effort was independently associated with breathlessness, a mixed effects regression model was created. Here, subjective breathlessness rating (1-10 Likert scale) was explained as a function of the physical effort of pedalling (W), VR hill gradient, state anxiety level before undertaking exercise (STAI questionnaire), age, sex, and BMI, with a random participant effect included. Following model optimisation through backwards elimination of non-significant model terms (see Supplementary Table 3 for the full and optimised models), physical effort of pedalling (W) (0.81 ± 0.21, 95% CI [0.39, 1.22], *p*<0.001, R^2^ = 0.175) and virtual hill gradient (0.63 ± 0.11, 95% CI [0.43, 0.84], *p*<0.001, R^2^ = 0.192) were positively associated with subjective breathlessness ratings, whilst BMI was significantly negatively associated (−0.72 ± 0.28, 95% CI [−1.28, −0.17], *p*=0.02, R^2^ = 0.198) (Figure 4). Variance inflation was low (<1.2) for all model terms, suggesting minimal collinearity.

**Figure 4.**
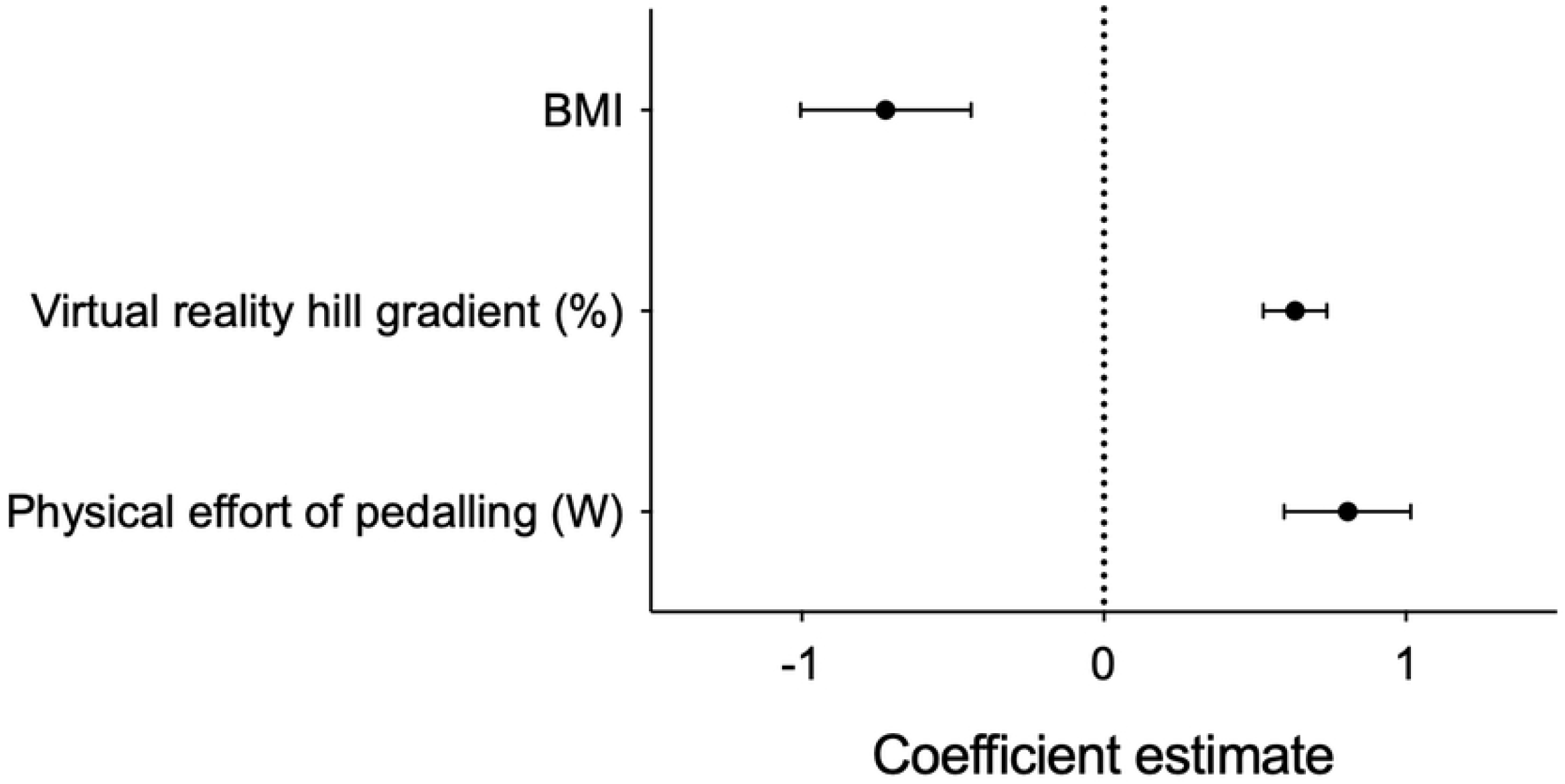
The physical effort of pedalling (actual effort), virtual hill gradient (expected effort), and BMI are independent predictors of an individual’s breathlessness.

**Figure 5.**
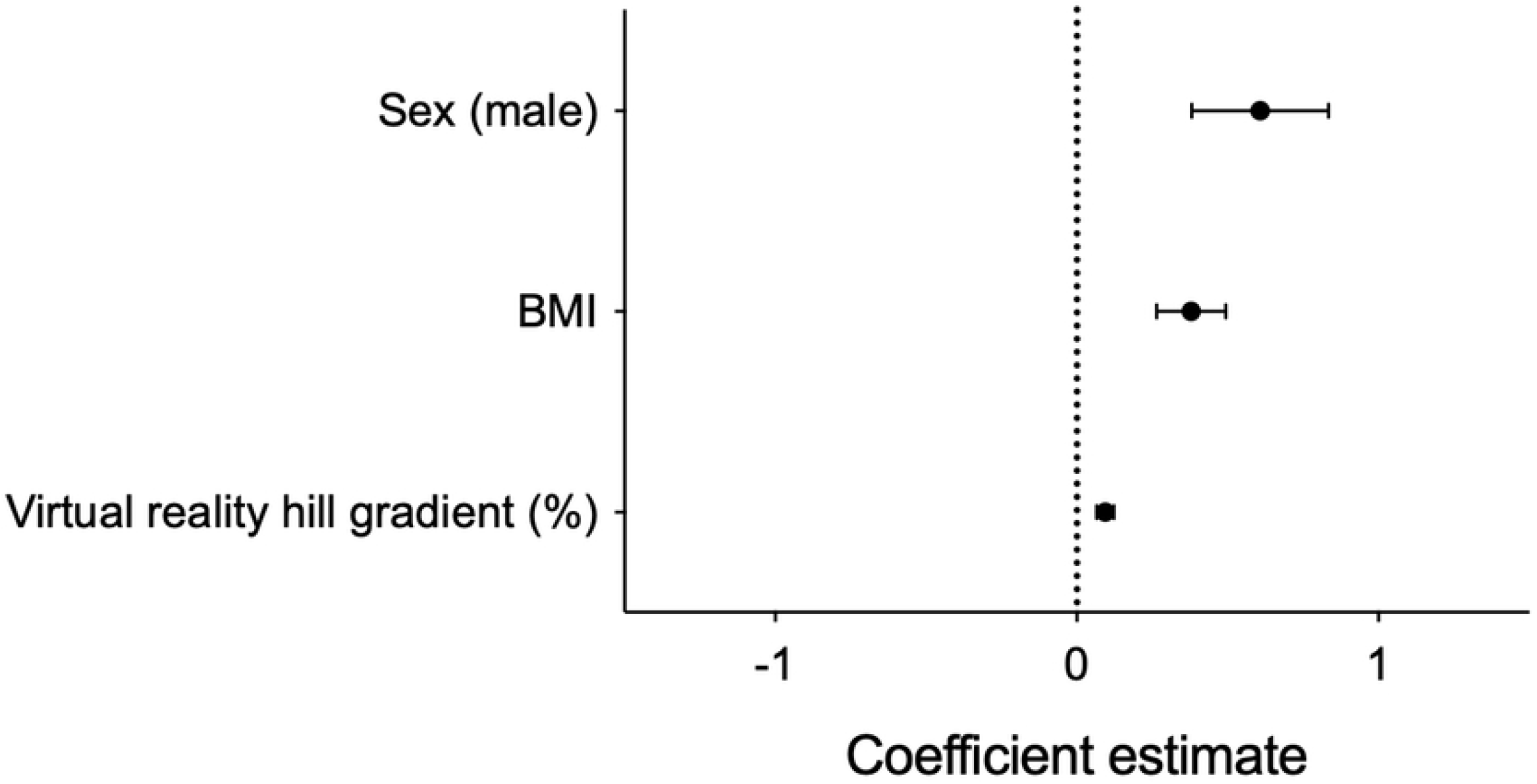
Virtual reality hill gradient, BMI, sex (male) are independent predictors of physical effort (i.e., the physical effort of pedalling)

### Effort expectation in VR modulates actual physical effort

Finally, we sought to determine whether effort expectation would influence participants’ actual effort. We created a second mixed effects model where the physical effort of pedalling (W) was explained as a function of virtual hill gradient, BMI, age, and sex, with a random participant effect included. Following model optimisation through backwards elimination of non-significant model terms (see Supplementary Table 4 for the full and optimised models), virtual hill gradient (0.09 ± 0.03, 95% CI [0.04, 0.15], *p*=0.001, R^2^ = 0.060), BMI (0.38 ± 0.11, 95% CI [0.15, 0.60], *p*=0.004, R^2^ = 0.518), and sex (male) (0.61 ± 0.23, 95% CI [0.16, 1.06], *p*=0.02, R^2^ = 0.411) were positively associated with the physical effort of pedalling. Variance inflation was low (1.0) for all model terms, suggesting minimal collinearity.

### Immersion in the VR cycling world

Each question on the presence questionnaire is scored out of a maximum of 7 points on a Likert scale, with one being “not at all” and seven being “very much”. Participants reported an average of 5.25 (IQR 1). Highest item scores were recorded for rapid adjustment to the virtual world and for the quality of display. Lowest scores were recorded for the item examining how consistent participants felt their experiences in the virtual world were compared to the real-world (Table 3).

**Table 3.** Item and total scores for the presence questionnaire (PQ).

| Question | Median score out of 7 (IQR) |
| --- | --- |
| How natural was the mechanism that controlled movement through the environment | 4.5 (1.5) |
| How compelling was your sense of objects moving through space? | 5 (2) |
| How much did your experiences in the virtual environment seem consistent with your real-world experiences? | 4 (1) |
| How involved were you in the virtual environment experience | 5.5 (1) |
| **How quickly did you adjust to the virtual environment experience** | 7 (1.5) |
| **How much did the visual display quality interfere or distract you from performing assigned tasks of required activities? (reverse scored)** | 6 (2.5) |
| **Total average score** | 5.25 (1) |

## Discussion

### Key findings

To form subjective sensations of breathlessness, the brain interprets afferent signals based on a set of held “expectations” [1–3]. In this study, we used a novel VR cycling paradigm to dissociate actual effort (the power required to pedal) from an individual’s effort expectation (observed VR hill gradients) and in doing so, determined the contribution of effort expectation to the sensation of breathlessness. We found that approximately 19% of the breathlessness experienced was explained by the VR hill gradient observed by participants, whereas approximately 18% was explained by the power required by participants to pedal.

Additionally, we showed that participants worked harder when they observed a steeper virtual slope, despite ergometer resistance being kept constant, with the observed VR hill gradient accounting for 6% of the power produced by participants. These findings demonstrate that an individual’s expectations of effort can independently modulate subjective perceptions, including breathlessness, and influence physical effort.

### Examining the contribution of expectations in chronic breathlessness

A growing body of evidence [1–3, 5, 9, 11, 25–28] now suggests that breathlessness, far from simply arising as a product of peripheral inputs, includes steps of active interpretation and comparison with previously held expectations. In this study we demonstrated that as the observed VR hill gradient increased, participants’ reported breathlessness was also increased, independent of the physical effort applied to pedalling the bike. This highlights that an individual’s prior expectations may influence their subjective experiences of exertion, including how breathless they feel in response to a given exercise intensity. This may help explain instances of unexplained breathlessness such as that observed in asthma or panic disorder, where breathlessness fails to match physiological assessments of cardiopulmonary health. This work is in line with previous studies showing, in cases of chronic breathlessness, that improvements in breathlessness are associated with a reappraisal of the sensory experience, i.e., changes to expectations [10] rather than increased fitness. However, determining the contribution of ‘expectation’ to the subjective human exercise experience is problematic, particularly where beliefs or expectations are difficult to articulate. Our paradigm may now provide a tool with which to unpick this relationship.

### Physical effort is influenced by virtual slope

In the real world, road inclinations are met with proportional increases in resistance due to gravitational force and because of learning about our physical environment, our expectation is that we must apply greater power to maintain our speed. In this experiment we observed a positive association between the virtual slope gradient and the pedalling power applied by participants, independent of ergometer resistance. To the authors knowledge, this is the first study to demonstrate that manipulating effort perception within a virtual environment influences an individual’s physical effort. There is a longstanding interest in whether hypnotic suggestion (perception manipulation) can modulate exercise capacity; however, results are equivocal [29]. While, our study did not investigate exercise performance, this finding nevertheless suggests that manipulating sensory or environmental cues could be used to increase physical capacity. Moreover, it is plausible that manipulation of sensory cues, as demonstrated here, could be used to increase physical effort in rehabilitation settings (e.g., cardiac recovery) and more broadly, training of the general population.

### Limitations

The purpose of this study was to demonstrate that subjective breathlessness ratings could be manipulated independently from physical work effect. Given the novelty of the design and lack of established evidence within the field to draw upon we have identified a number of key limitations that future studies will work to address.

In terms of hardware, while VR headsets provide a fully immersive experience, the headset generates a noticeable amount of heat, which paired with physical exertion can quickly become uncomfortable for the participant. To overcome this, future studies may investigate other display options such a wide screen monitors or enlist newer headsets with built in cooling systems. In this single session we assessed the extent to which the illusion was maintained across a wide range of virtual slopes. However, the Tacx turbo trainer was only capable of simulating gradients of 0% to 6%. Consequently, the range of physical slopes did not align with the range of virtual slopes. Future studies would improve statistical power by limiting the range of virtual slopes under comparison and extend either the number of sessions or time over which data was collected. Additional psychological and physiological characterisation would also help to answer some outstanding questions. Further physiological measures, quantified by cardiopulmonary exercise testing would help to answer questions regarding whether the virtual environment simply changed perceptions of work effort on a cognitive level or whether an individual’s physiology was also affected. Previous work using hypnosis has demonstrated that manipulating effort perception activates cortical structures involved in the cardiopulmonary response to exercise, resulting in increased blood pressure and heart rate [15]. While in terms of psychological characterisation the presence questionnaire is somewhat limited by a lack of validated thresholds. Further assessment of anxiety and interoceptive sensitivity and its relationship with manipulability of expectation may also be of interest. Studying more highly anxious individuals would be worthwhile as we might predict that the effects of VR manipulation may be even greater. Finally, this study has provided initial evidence for the manipulation of breathlessness expectations within a healthy population. Further work would extend this paradigm to assess utility in clinical populations.

### Future directions

#### Realigning expectations in chronic breathlessness

Realigning expectations with sensory inputs and challenging aberrant thought processes is a key foundation of many cognitive behavioural therapies. Compelling evidence highlights how pulmonary rehabilitation, the current most effective treatment for chronic breathlessness in COPD, has little effect on lung function but significantly improves feelings of breathlessness via changes in brain processes [10]. While a number of brain-targeted therapeutic approaches, including pharmacological [30, 31], direct brain stimulation [32] and cognitive [33] have been explored with varying degrees of success, VR offers another approach in which sensory input may be realigned with expectation in an example of “fool the brain, treat the lungs” [11]. The high entertainment value of immersive VR and increasing affordability also has clear potential within a framework of pulmonary rehabilitation to promote exercise engagement and expose patients to “real-life” environments in which to overcome fearful breathlessness safely.

### Conclusions

Using a multi-sensory, fully immersive, VR cycling environment we have shown that perceived breathlessness can be driven independently from physical work effort. This finding highlights the importance of expectation in breathlessness perception which have potential implications for training and rehabilitation programmes.

## Acknowledgements

The authors would like to thank Silvia Pan for her valuable input into the manuscript.

